# Polystyrene and polyethylene terephthalate nanoplastics differentially impact mouse ovarian follicle function

**DOI:** 10.1101/2025.06.27.662011

**Authors:** Hanin Alahmadi, Maira Nadeem, Alixs M. Pujols, Raulle Reynolds, Mohammad Saiful Islam, Indrani Gupta, Courtney Potts, Allison Harbolic, Gania Lafontant, Somenath Mitra, Genoa R. Warner

## Abstract

Exposure to micro- and nanoplastics is unavoidable. Foods and beverages contain plastic particles from environmental contamination and processing and packaging materials, which are frequently made of polyethylene terephthalate (PET). Micro- and nanoplastics have been detected in human tissues such as the brain, liver, and placenta, as well as in ovarian follicular fluid, but little is known about the effects nanoplastics have on the female reproductive system. In addition, few studies on the health impacts of nanoplastics have been performed using environmentally relevant plastic types and concentrations. Thus, this research tested the hypothesis that nanoplastics made of spherical polystyrene (PS), a common model nanoplastic, would have different effects on cultured mouse ovarian follicles compared to secondary PET nanoplastics at environmentally relevant doses. The ovary is a highly sensitive reproductive organ responsible for the development of follicles, which contain the oocyte, and production of steroid hormones. Follicles were harvested from adult mouse ovaries and cultured for 96 h with vehicle, spherical commercially available 200 nm PS nanoplastics (1–100 µg/mL), or lab-generated 240 nm PET nanoplastics (0.1–10 µg/mL). PS and PET nanoplastic exposure inhibited follicle growth and altered expression of genes related to steroid synthesis, cell cycle, and oxidative stress. PET nanoplastics increased levels of pregnenolone and decreased expression of *Cyp17a1*. Overall, both plastic types altered ovarian function, but they impacted different genes in similar pathways. These findings suggest that nanoplastic exposure at environmentally relevant concentrations may pose a risk to female reproductive health by disrupting hormonal and molecular pathways. In addition, environmentally relevant plastic types and doses are necessary for studying health impacts of nanoplastics.

**Highlights:**

- Nanoplastics disrupted ovarian function in vitro
- Secondary PET and virgin PS differently inhibited follicle growth and ovarian gene expression
- PET exposure increased pregnenolone, impacting steroid hormone pathways
- Environmentally relevant plastics can harm female reproductive health

## 1 Introduction

Plastics are widely popular due to their low cost and versatility, which has led to increasing production over the past century. Due to this, the term "Age of Plastics" is used to describe the modern era (Avio et al., 2017). In 2022, global annual plastic production was estimated to reach over 400 million tons and continues to exponentially increase (Nayanathara Thathsarani Pilapitiya and Ratnayake, 2024). Common plastic polymers were historically thought to be inert, and are widely used in food, beverage, and personal care product packaging and medical equipment. However, studies suggest that plastic materials shed micro- (<5 mm) and nano-sized (<1 μm) plastic particles during use, such as into drinking water (Mason et al., 2018; Qian et al., 2024). In addition, plastic particles generated by weathering of environmental plastic waste contaminate our food, water, and air (Giri et al., 2024). Both bottled and tap water have been found to contain plastic particles, as well as foods with trophic accumulation such as seafood (Senathirajah et al., 2021). Inhalation is also a significant source of exposure from synthetic microfibers and urban dust (Giri et al., 2024). Intake estimates are uncertain and vary widely (Zarus et al., 2021). One widely publicized estimate of human exposure from all sources calculated 0.1 – 5 g per person per week, which is equivalent to about 0.2–9 mg/kg/day in an adult woman (Senathirajah et al., 2021). In another study, daily intake of microplastics from bottled water was estimated at 40 μg/kg/day (Zuccarello et al., 2019). Recent studies of human feces and indoor dust estimated intake of polyethylene terephthalate (PET), a common polymer used in food and beverage packaging, at 83–120 μg/kg/day for infants and 5.8–6.6 μg/kg/day for adults (Zhang et al., 2021, 2020).

Studies identifying plastic particles in human tissues reveal that the particles remain in our bodies. Strikingly, recent analyses of human blood, placentas, testes, livers, kidneys, brains, and other tissues confirm that nanoplastics as well as larger plastic particles are universally present (Brits et al., 2024; Garcia et al., 2024; Hu et al., 2024; Leslie et al., 2022; Nihart et al., 2025). Most of these studies on human organs contain an average of 100–400 μg/g plastic, as measured by pyrolysis gas chromatography mass spectrometry, although brains contained about ten times more. Estimates in human blood include averages of 1.6 μg/mL (Leslie et al., 2022) and 0.27 μg/mL (Brits et al., 2024) with maximum samples values of 12 and 2.5 μg/mL, respectively. Microplastics were recently reported in human ovarian follicular fluid (Montano et al., 2025).

Despite confirmation that tiny plastic particles are universally present in tissues, including sensitive reproductive organs, little is known about the health implications. The ovary is the primary site of steroid hormone production and fertility, and is sensitive to disruption by environmental contaminants, especially endocrine disruptors (Patel et al., 2015). *In vitro* and animal studies suggest that plastic particles can disrupt fertility, hormone levels, and/or ovarian function in oysters (Sussarellu et al., 2016), medaka fish (González-Doncel et al., 2022), zebrafish (Limonta et al., 2019), mice (Huang et al., 2023; Liu et al., 2022; Wang et al., 2025; Wei et al., 2022; Xiong et al., 2024; Xue et al., 2024; Zeng et al., 2023; Zhang et al., 2023), rats (Haddadi et al., 2022; Marcelino et al., 2022), and cultured human granulosa cells (Huang et al., 2023; Xue et al., 2024; Zeng et al., 2023; Zhang et al., 2025). For example, exposure to 33 mg/kg of 25 nm nanoplastics for 42 days disrupted steroidogenesis, interfered with hormone production, and impaired oocyte quality in mice, potentially leading to reduced fertility in females (Xue et al., 2024). In addition, direct comparison of male and female mice suggests that females may be more sensitive (Wei et al., 2022). Together, these studies suggest that nanoplastics pose a risk for ovarian function. However, nearly all previous studies used high doses that are not environmentally relevant and spherical virgin polystyrene (PS) beads to model plastic exposure, which are not representative of environmental exposure to weathered and secondary nanoplastics. The effects across plastic polymer types, shapes, and sizes are still not well understood.

In this study, we compare the impacts of nanoplastics made of two common polymers, PS and PET, on ovarian function, specifically folliculogenesis, the development of the oocyte-containing ovarian follicle, and steroidogenesis, the production of sex steroid hormones. We utilized an *ex vivo* mouse follicle culture system to test the hypothesis that PS and PET nanoplastics harm ovarian function at environmentally relevant doses and that effects vary with particle properties. This knowledge is important because disruption of ovarian processes poses risks to fertility and hormone balance.

## 2 Method and Materials

### 2.1 Nanoplastics

200 nm polystyrene (PS) nanospheres were acquired from ThermoFisher Scientific (R200). Polyethylene (PET) nanoplastics were ground in the laboratory from a plastic milk bottle using a method described previously (Islam et al., 2024). PS was chosen because of its commercial availability and to compare to previous studies. PET was chosen because it is a common polymer type found in the environment and human tissues and because it is used to bottle drinking water (Li et al., 2024). Particles were characterized using scanning electron microscopy (SEM), dynamic light scattering (DLS), and Fourier transform infrared spectroscopy (FTIR).

### 2.2 Animals

Young adult female CD-1 mice (23 days old) were purchased from Charles River (Wilmington, MA). They were acclimatized in a controlled environment at the Rutgers University Newark Animal Facility for at least 7 days, with a 12-hour light-dark cycle, 22 ± 1 °C temperature, and ad libitum access to food and water. All experiments were performed with Rutgers Institutional Animal Care and Use Committee (IACUC) approval (PROTO202100043).

### 2.3 Antral follicle culture

For the follicle culture studies, adult CD-1 mice of age 30-38 days were euthanized. Antral follicles of sizes 200 to 400 µM were mechanically harvested from the ovaries. Follicles were cleaned from interstitial tissues using watchmaker forceps. For each replicate, follicles collected from 4-5 mice were individually cultured in 96 well plates with 10 to 12 follicles per treatment group. Treatment groups were either vehicle controls, PS nanoplastics (1 – 100 µg/mL), or PET nanoplastics (0.1 – 10 µg/mL). Each treatment group was combined with supplemented media consisting of minimum essential medium alpha (Gibco) containing 1 % insulin-transferrin-selenium (10 ng/mL insulin, 5.5 mg/mL transferrin, 5 ng/mL sodium selenite, Sigma-Aldrich), 100 U/mL penicillin (Sigma-Aldrich), 100 mg/mL streptomycin (Sigma-Aldrich), 5 IU/mL human recombinant follicle-stimulating hormone (Scripps Laboratories, San Diego, CA), and 5 % fetal bovine serum (Gibco).

Media were treated with appropriate NP concentrations and were not changed during culture. Stock concentrations were prepared for PS nanoplastics (1 – 100 µg/mL) in 5% Tween20 and PET nanoplastics (0.1 – 10 µg/mL) in cell culture grade water (Corning) and stored in the fridge. 10 µL of stock was used per 1 mL of supplemented media for a final Tween20 concentration of 0.05% for PS. Different vehicles were used to match the solutions that the particles were provided in, and each plastic type was compared to its respective control. There were no statistically significant differences in follicle growth between the two vehicle control groups. Follicles were cultured for 96 hours in an incubator with 5% CO_2_ supply and 37 °C.

### 2.4 Follicle growth analysis

The growth of cultured antral follicles over 96 h was assessed at 24 h timepoints by measuring follicle diameters on perpendicular axes with an inverted microscope equipped with a calibrated ocular micrometer. The diameters of each follicle were averaged within the treatment group for each 24 h interval, and the average values were divided by the initial average measurement at 0 h of each of the respective treatment groups to calculate the percent change in follicle diameter over time. The percent change in antral follicle diameter over time was used for statistical analysis.

### 2.5 Hormone assays

After 96 hours of culture, media from each treatment group were combined and stored at −80 °C until analysis. Enzyme-linked immunosorbent assays (ELISAs, DRG International Inc., Springfield, New Jersey) were used to measure levels of pregnenolone, progesterone, testosterone, estradiol, and pregnenolone. Intraassay and interassay coefficients of variability were, respectively: pregnenolone 14.3% and 17.6%; progesterone 13.7% and 14.4%; androstenedione 3.5% and 11.2 %; testosterone 8.5% and 2.6%; estradiol 10.7% and 13.7%.

### 2.6 Gene expression analysis

After culture, follicles from each treatment group were combined and stored at −80 °C until analyses. Total RNA was extracted from the follicles using either the E.Z.N.A. MicroElute® Total RNA Kit (Omega Bio-tek, Norcross, Georgia), or the Qiagen RNeasy Micro Kit (Qiagen, Germantown, MD), according to the respective manufacturer’s protocol. The RNA concentrations were quantified using Thermo Scientific Nanodrop One C Spectrophotometer. To create cDNA, the RNA was reverse transcribed using iScript Reverse Transcription Kit (Bio-Rad Laboratories, Inc., Hercules, California). Quantitative PCR (qPCR) was conducted using the Bio-Rad CFX Duet PCR instrument to analyze the gene expression levels. The qPCR was conducted with the following cycling conditions: 94°C for 30s, 42 cycles at 94°C for 5s, and 60 °C for 34s. Reactions were run in duplicate with 1.67 ng cDNA and 7.5 pmol of gene-specific primers (Integrated DNA Technologies, Inc. Coralville, Iowa) for a final reaction volume of 10 µl. Genes analyzed include cell cycle regulators cyclin A2 (*Ccna2*), cyclin B1 (*Ccnb1*), cyclin-dependent kinase 4 (*Cdk4*), cyclin-dependent kinase inhibitor 1A (*Cdkn1a*), cyclin-dependent kinase inhibitor 1B (*Cdkn1b*); apoptosis related genes B cell leukemia/lymphoma 2 (*Bcl2*), Bcl2-associated X protein (*Bax*), BCL2-associated agonist of cell death (*Bad*), caspase-8 (*Casp 8*), caspase 3 (*Casp3*); oxidative stress related genes NADPH oxidizer organizer 1 (*Noxo1*), superoxide dismutase 1 (*Sod1*), and glutathione peroxidase 1 (*Gpx1*); hormone receptor related genes estrogen receptor 1 (*Esr1*), estrogen receptor 2 (*Esr2*), follicle stimulating hormone receptor (*Fshr*) and androgen receptor (*Ar*); steroidogenesis related genes steroidogenic acute regulatory protein (*StAR*), cholesterol side-chain cleavage enzyme (*Cyp11a1*), cytochrome P450 family 17 subfamily A member 1 (*Cyp17a1*), hydroxy-delta-5-steroid dehydrogenase, 3 beta- and steroid delta-isomerase 1 (*Hsd3b1*), hydroxysteroid 17-beta dehydrogenase 1 (*Hsd17b1*), aromatase (*Cyp19a1*) and compared between treatment groups. qPCR analysis was done using the Pfaffl method with the housekeeping gene beta-actin (*ActB*) as the reference gene (Pfaffl, 2001). Genes are listed in **Table 1**.

**Table 1:**
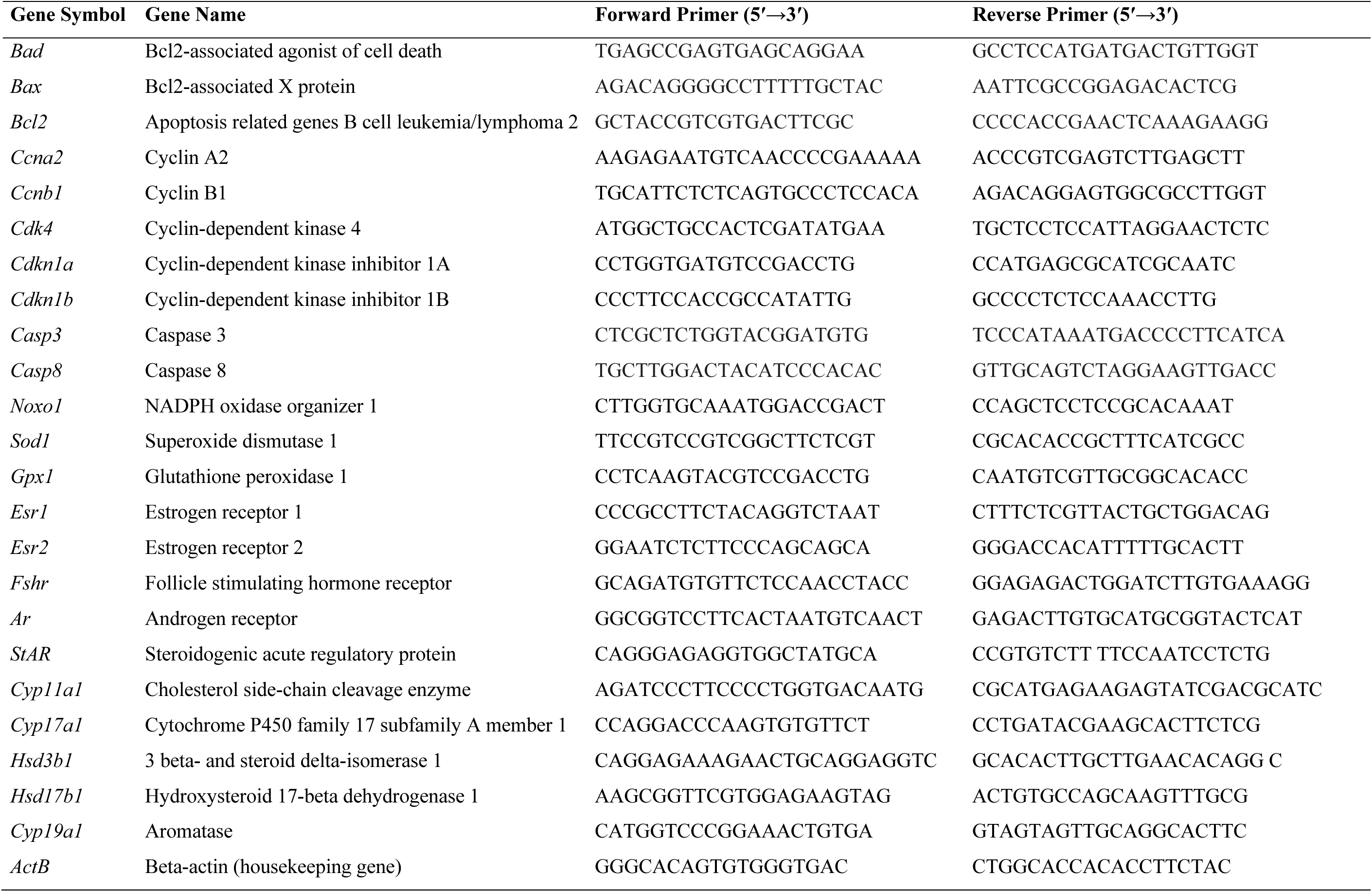
DNA oligonucleotide primers used for qPCR gene expression assays.

### 2.7 Statistical analysis

Data are expressed as means ± standard error of the mean (SEM) from at least 3 separate experiments. Data were analyzed by comparing treatment group to control using IBM SPSS version 29 software (SPSS Inc., Chicago, IL, USA). All data were continuous and assessed for normal distribution by Shapiro-Wilk analysis. If data met assumptions of normal distribution and homogeneity of variance, data were analyzed by one-way analysis of variance (ANOVA) followed by Tukey HSD or Dunnett 2-sided post-hoc comparisons. However, if data met assumptions of normal distributions, but not homogeneity of variance, data were analyzed by ANOVA followed by Games-Howell or Dunnett’s T3 post-hoc comparisons. If data were not normally distributed or presented as percentages, the independent sample Kruskal-Wallis H followed by Mann-Whitney U non-parametric tests were performed. For all comparisons, statistical significance was determined by p-value ≤ 0.05. If p-values were greater than 0.05, but less than or equal to 0.10, data were considered to exhibit a trend towards significance.

## 3 Results

### 3.1 Particle Characterization

Nanoplastic particles used in tissue culture experiments were characterized by SEM, DLS, and FTIR. Characterization confirmed the average particle size of purchased PS nanoplastics at 220 nm with a polydispersity index (PDI) of 0.085 (**Figure 1A**). Secondary PET nanoplastics generated in the laboratory were found to agglomerate in solution. Before sonication, the average diameter was 661 nm with PDI of 0.047, whereas after sonication the particles were 240 nm with PDI of 0.462 (**Figure 1B,C**). Signature FTIR peaks of PS were 3024 cm^-1^, for aromatic C–H stretches, 2918 and 2848 cm^-1^ for CH_2_ stretches, 1592 and 1451 cm^-1^ for aromatic C–C stretches, and 758 and 696 cm^-1^ for aromatic C-H bends (**Figure S1**). The PET exhibited distinctive peaks at 2923 and 2853 cm^-1^ for C–H stretches, 1718 cm^-1^ for the C=O ester group, 1410 cm^-1^ for –CH2– deformation band, 1240 and 1096 cm^-1^ for C–C–O and C–O–O stretching of ester groups, respectively, and 727 cm^-1^ for C–H out-of-plane deformation of two carbonyl substituents on the aromatic ring (**Figure S1**)

**Figure 1.**
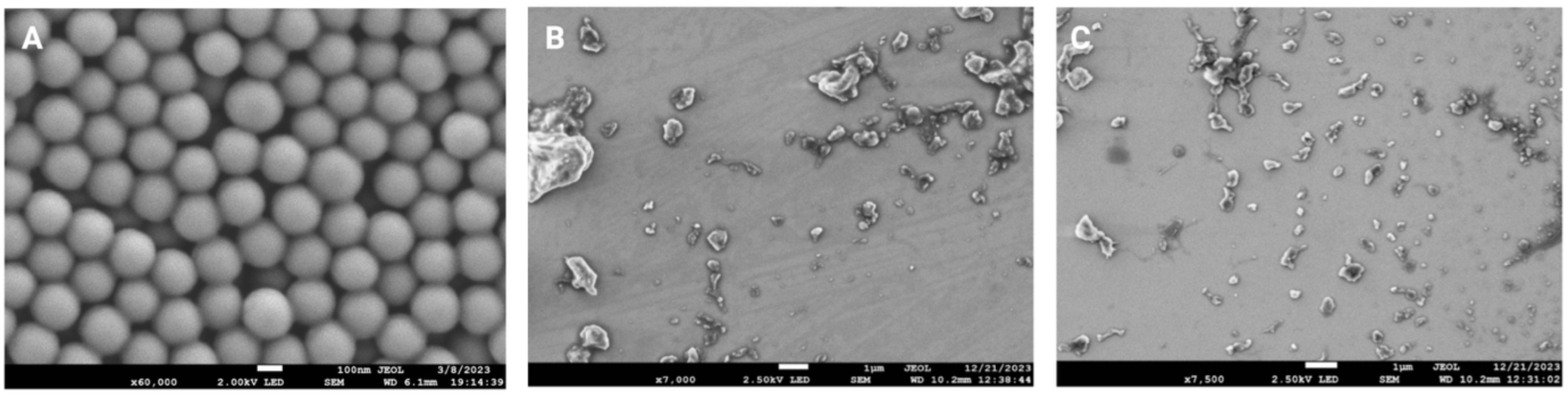
Nanoplastics were characterized using scanning electron microscopy (SEM) for purchased PS spheres (A, scale bar 100 nm) and lab ground PET before (B) and after (C) sonicating (scale bar 1µm).

### 3.2 Effects of nanoplastic exposure on follicle growth

Antral follicles isolated from mouse ovaries were cultured in media containing vehicle control, PS nanoplastics (1 – 100 µg/mL), or PET nanoplastics (0.1 – 10 µg/mL) for 96 hr (**Figure 2**). Follicle growth was inhibited in the 100 μg/mL PS treatment group at 48 hours (p = 0.047) and 72 hours (p = 0.076) compared to control. Follicle growth was also inhibited from 1 μg/mL PET (p = 0.076) and 10 μg/mL PET (p = 0.009) at 96 hours compared to control.

**Figure 2.**
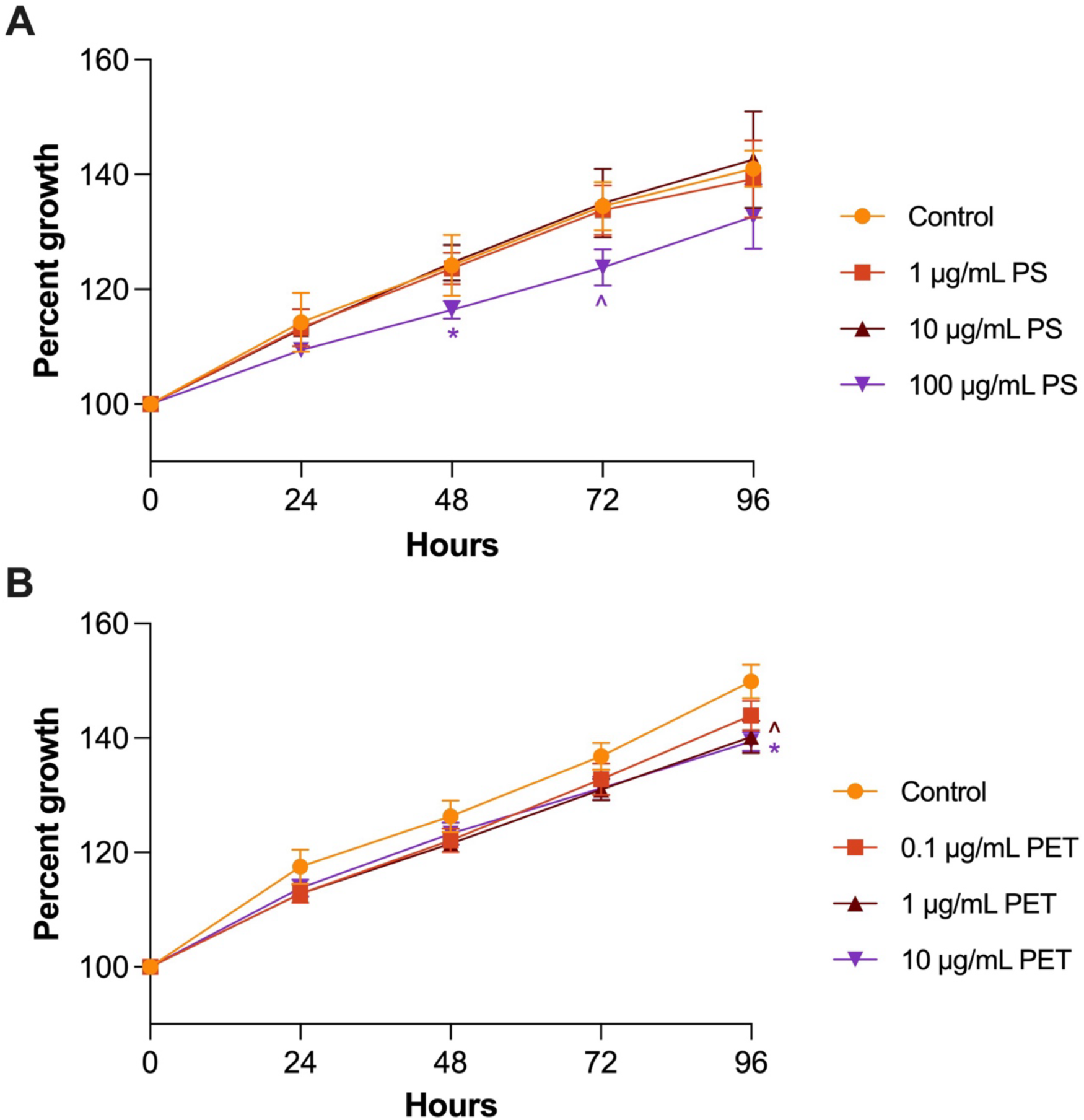
Effect of PS (A) and PET (B) treatment on antral follicle growth. Follicle growth was measured every 24 hrs for 96 hrs. Graphs represent mean ± SEM from 5–6 independent replicates per treatment group. Asterisks (∗) indicate significant differences from the control (p ≤ 0.05) and ^ indicates a trend toward significance.

### 3.4 Effects of nanoplastic exposure on sex steroid hormone levels

PS nanoplastic exposure did not significantly alter steroid hormone levels compared to control (**Figure 3A**). Pregnenolone was significantly increased at 10 μg/mL PET and 100 μg/mL PET compared to control (**Figure 3B**).

**Figure 3.**
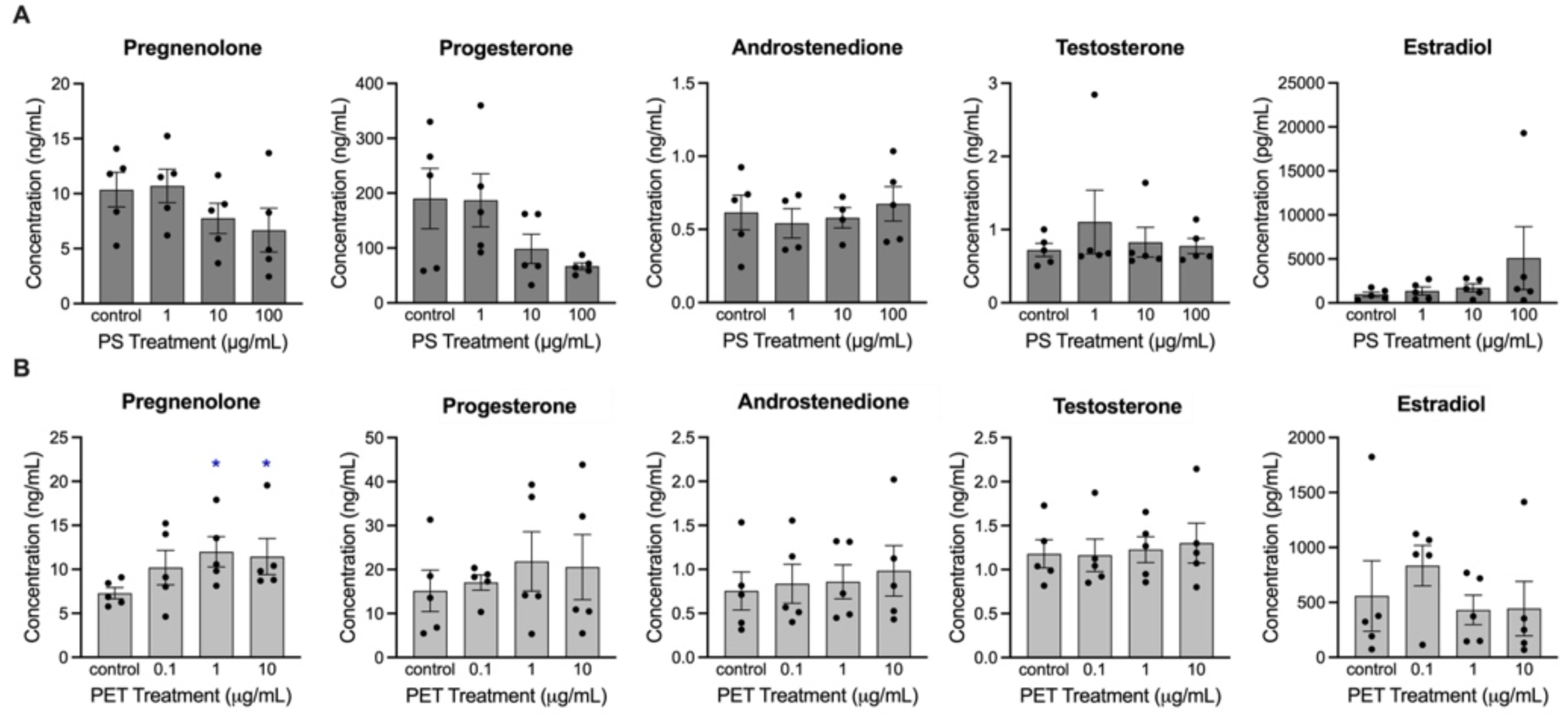
Impact of PS (A) and PET (B) treatment on hormone levels in the media following 96 hrs of culture. Culture media were subjected to enzyme-linked immunosorbent assays. Asterisks (∗) indicate significant differences from the control (p ≤ 0.05). Graphs represent mean ± SEM from 4–6 separate experiments. Asterisks (∗) indicate significant differences from the control (p ≤ 0.05) and ^ indicates a trend toward significance.

### 3.5 Effects of nanoplastic exposure on gene expression in antral follicles

#### 3.5.1 Steroidogenesis enzymes

Exposure to PS nanoplastics resulted in alterations in the expression of steroidogenesis genes (**Figure 4**). *StAR* expression was significantly decreased in the 100 μg/mL PS treatment group whereas *Cyp19a1* showed a borderline increase at the 10 μg/mL PS treatment group compared to the control. *Cyp17a1* was significantly decreased at 100 μg/mL PET compared to control

**Figure 4.**
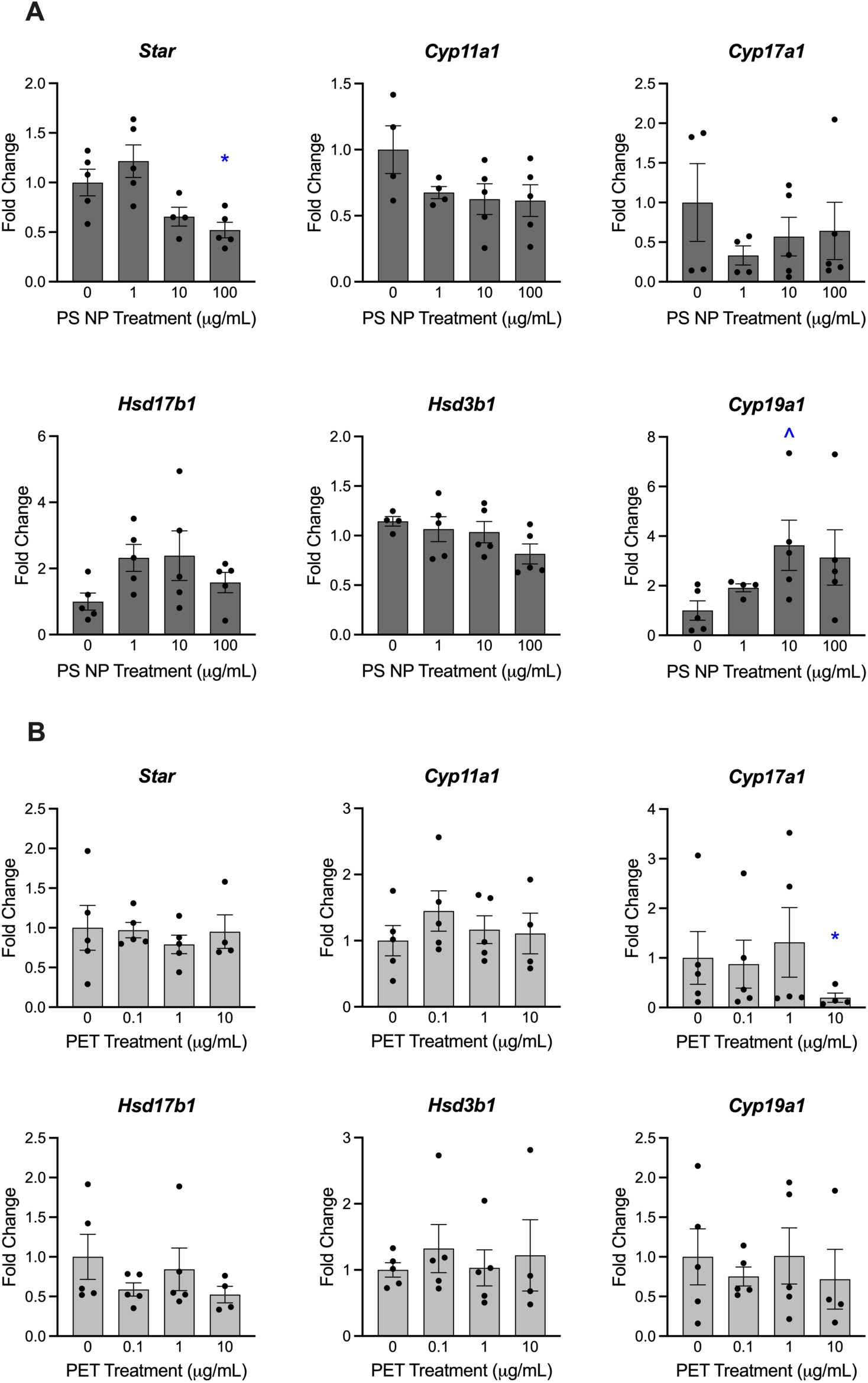
Impact of PS (A) and PET (B) treatment on steroidogenesis gene expression measured via qPCR. Graphs represent mean ± SEM from 4–6 separate experiments. Asterisks (∗) indicate significant differences from the control (p ≤ 0.05) and ^ indicates a trend toward significance (p ≤ 0.10).

#### 3.5.2 Cell cycle regulators

Exposure to PS nanoplastics altered expression of cell cycle regulators (**Figure 5**).

**Figure 5.**
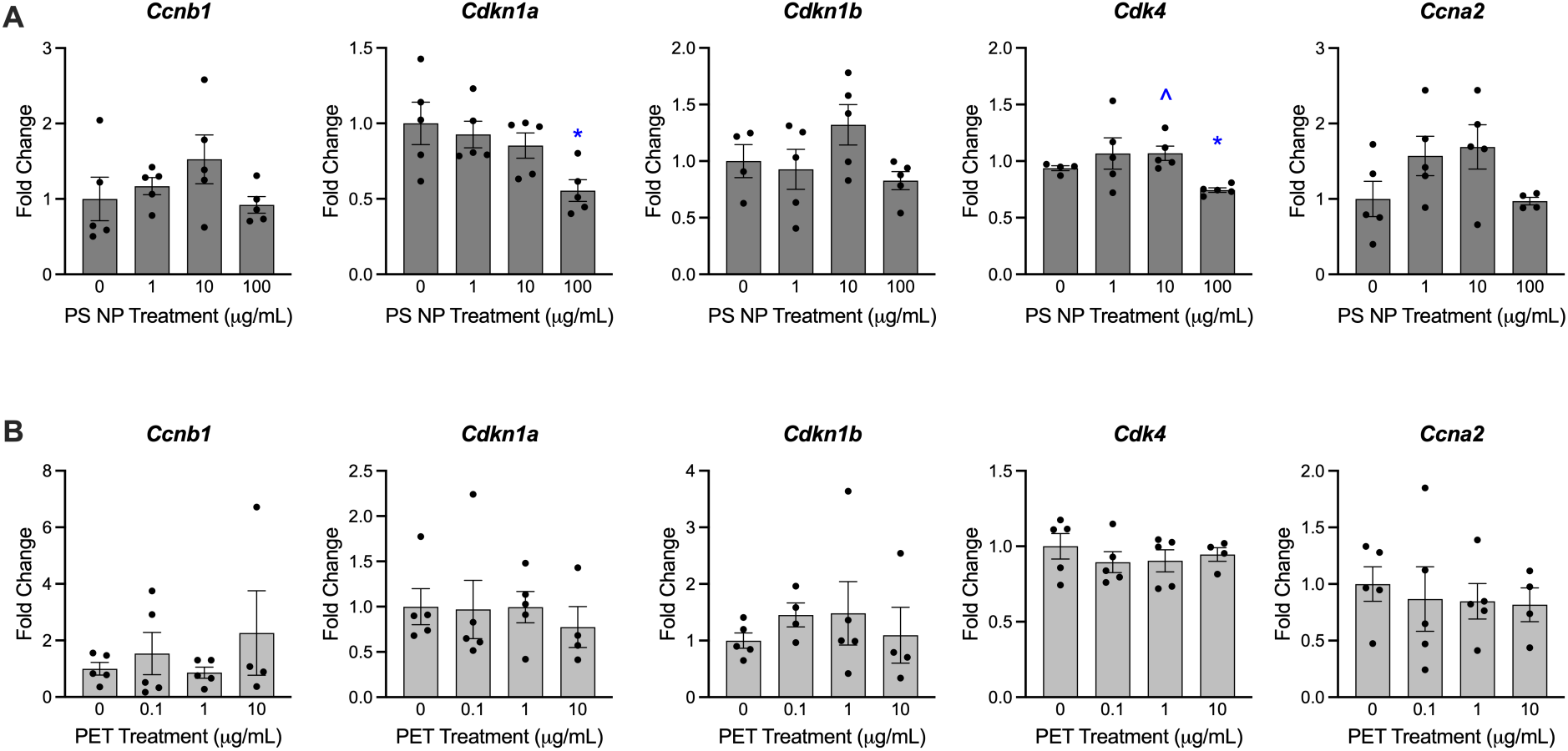
Impact of PS (A) and PET (B) treatment on cell cycle regulator expression measured via qPCR. Graphs represent mean ± SEM from 4–6 separate experiments. Asterisks (∗) indicate significant differences from the control (p ≤ 0.05) and ^ indicates a trend toward significance (p ≤ 0.10).

*Cdkn1a* expression was significantly decreased at 100 μg/mL PS compared to the control. *Cdk4* expression borderline increased at 10 μg/mL and significantly decreased at 100 μg/mL PS compared to control. No statistically significant impacts were observed from PET exposure.

#### 3.5.3 Apoptosis regulators

The apoptosis related gene *Bax* was significantly decreased at 100 μg/mL compared to the control. *Casp3* was significantly increased at 10 μg/mL PS and significantly decreased at 100 μg/mL PS compared to control (**Figure 6**). No statistically significant impacts were observed from PET exposure.

**Figure 6.**
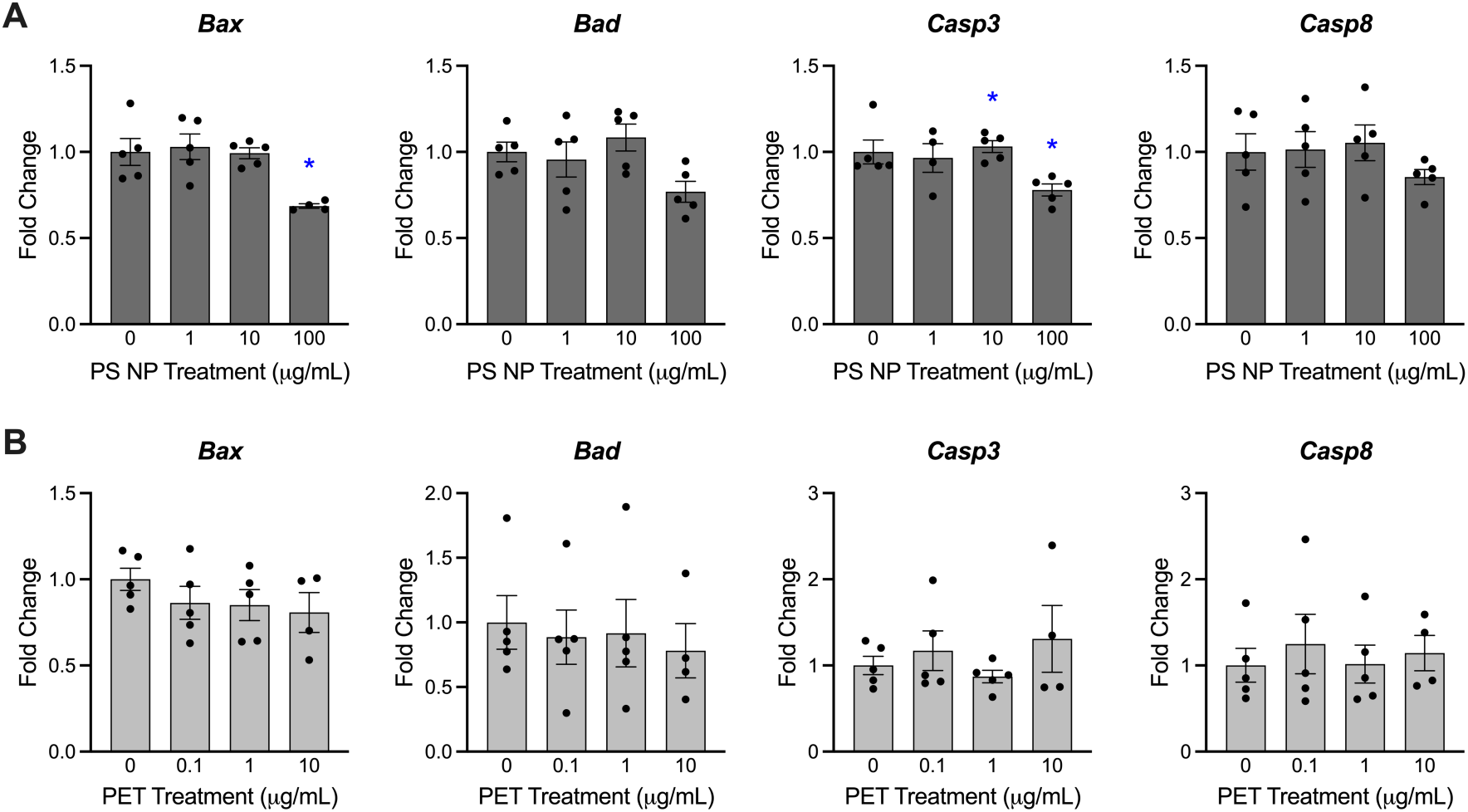
Impact of PS (A) and PET (B) treatment on apoptosis related gene expression measured via qPCR. Graphs represent mean ± SEM from 4–6 separate experiments. Asterisks (∗) indicate significant differences from the control (p ≤ 0.05) and ^ indicates a trend toward significance (p ≤ 0.10).

#### 3.5.4 Receptors

The expression of the androgen receptor gene *Ar* was significantly reduced at 100 μg/mL PS compared to the control (**Figure 7**). No statistically significant impact was observed from PET exposure.

**Figure 7.**
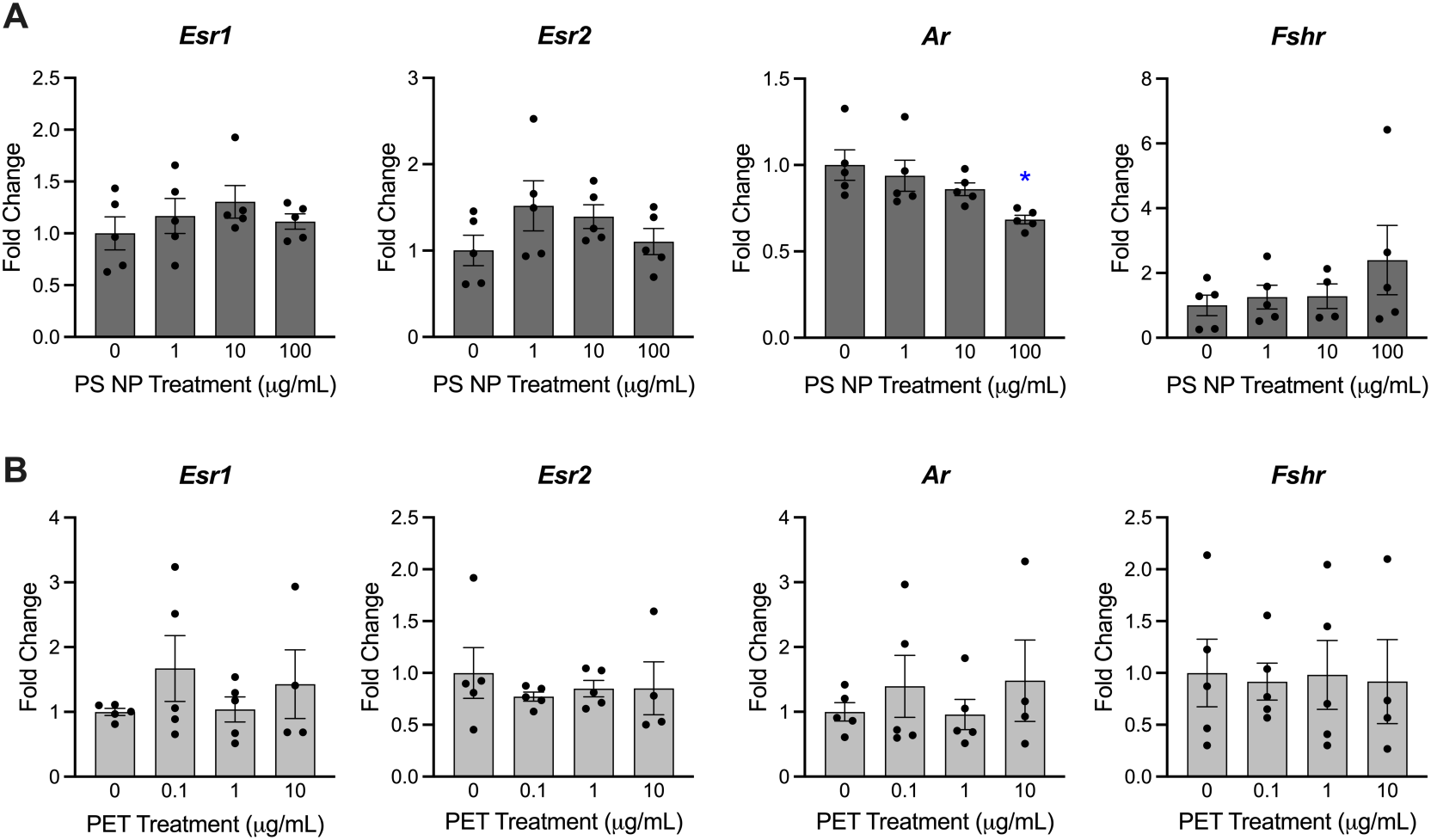
Impact of PS (A) and PET (B) treatment on receptor gene expression measured via qPCR. Graphs represent mean ± SEM from 4–6 separate experiments. Asterisks (∗) indicate significant differences from the control (p ≤ 0.05).

#### 3.5.5 Oxidative stress

The antioxidant gene *Sod1* exhibited a significant decrease at 100 μg/mL PS compared to the control (**Figure 8**). No statistically significant impact was observed from PET exposure.

**Figure 8.**
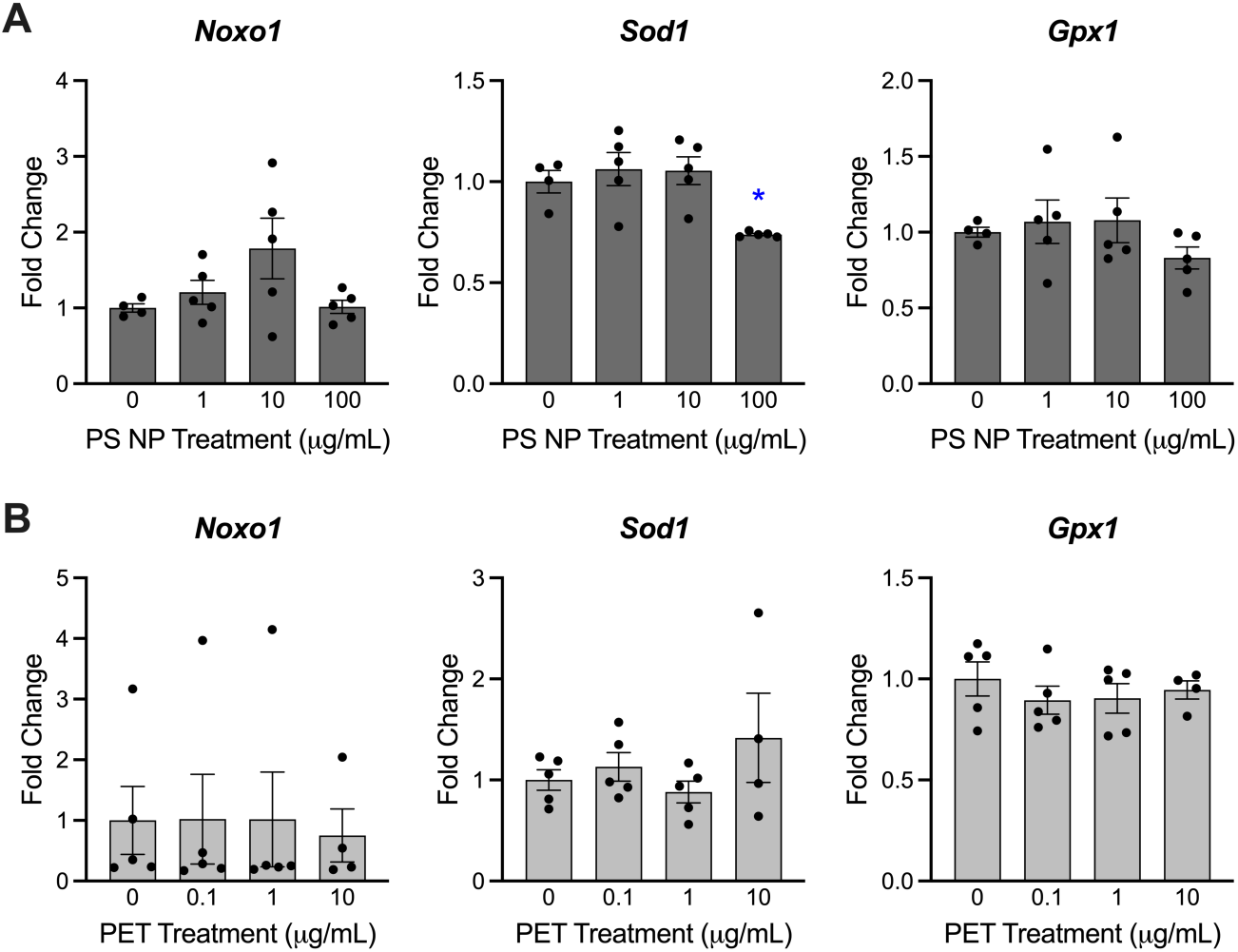
Impact of PS (A) and PET (B) treatment on oxidative stress gene expression measured via qPCR. Graphs represent mean ± SEM from 4–6 separate experiments. Asterisks (∗) indicate significant differences from the control (p ≤ 0.05).

## 4 Discussion

The adverse effects of nanoplastic exposure on hormones and reproductive function are not well understood. This gap in knowledge served as the primary motivation for conducting this study. Previous studies have shown effects from microplastics on the female reproductive system, such as ovarian dysfunction, granulosa cell apoptosis, and hormonal imbalance (Balali et al., 2024),(Afreen et al., 2023),(Yang et al., n.d.). However, few studies have been conducted on nanoplastics, and the few existing studies have focused on polystyrene spheres < 100 nm. We identified a gap in the literature looking at particles 100 – 1000 nm and of shapes, sizes, and polymers other than virgin PS spheres. Thus, we tested the hypothesis that secondary PET nanoplastics at environmentally relevant doses would disrupt ovarian function differently from PS spheres. Overall, we found that follicle growth, steroidogenesis, and expression of genes related to essential ovarian functions were differentially impacted by PS and PET.

PET is one of the most common plastic types and is widely used in food and beverage packaging. PET is one of the top five polymers detected in drinking and surface waters (Koelmans et al., 2019) and in the human body (Garcia et al., 2024). Not only does PET plastic waste weather and degrade into nanoplastics in the environment, but studies also show that PET bottles shed secondary particles during use, especially if single-use bottles are reused (Winkler et al., 2019). In a study comparing three brands of bottled drinking water, PET micro and nanoplastics were the most commonly detected polymer type in one brand of water and the second-most detected type in the other two (Qian et al., 2024), suggesting that humans are widely exposed to un- or lightly weathered PET particles. PET has been detected in human blood with average concentrations of 0.27–1.6 μg/mL and maximum sample value of 12 μg/mL (Brits et al., 2024; Leslie et al., 2022). Thus, we used doses of 0.1 – 10 μg/mL for PET to represent internal levels. Previous studies with PET micro and nanoplastics *in vitro* and in mice and zebrafish found increased oxidative stress, genotoxicity, alterations in gut microbial composition, and disturbances in metabolic pathways such as lipid regulation (Annangi et al., 2023; Bashirova et al., 2023; Ji et al., 2020; Lin et al., 2023; Manoochehri et al., 2024; Zhang et al., 2022), but no previous studies have looked at the impacts of PET particles on female reproduction or the ovary.

In our study, nanoplastic exposure led to significant changes in hormone levels and genes coding for steroidogenic enzymes early in the steroidogenesis pathway. The most noticeable impact was on pregnenolone levels, which increased at 1 and 10 μg/mL PET compared to control, and *Cyp17a1*, which codes for CYP17A1 to convert pregnenolone to dehydroepiandrosterone and progesterone into androstenedione, was significantly decreased at the highest treatment group. These opposing effects suggest a compensatory response to account for disruption, although it is unclear which is the chicken and which is the egg. PS exposure also impacted early steroidogenesis, with *Star* downregulated at the highest dose. This is the first study to directly expose whole antral follicles to nanoplastics and to investigate the direct impacts on the follicle. No previous in vitro ovarian cell studies have investigated steroidogenesis, but in vivo rodent studies have found decreased expression of *Star, Cyp11a1*, and *Cyp19a1* following 42 days of 1 mg/kg/day exposure to 25 nm PS nanoplastics in adults (Xue et al., 2024) and decreased serum progesterone levels with no impact on estradiol from 250-270 μg/kg/d exposure to 95 nm PS for 28 days. Conversely, other studies have shown that nanoplastics or microplastics exposure decreases serum estradiol levels (these studies did not measure progesterone) (Huang et al., 2023; Wei et al., 2022; Xiong et al., 2024). We may not have observed this effect because in vivo studies capture brain-gonad signaling or because of our small sample size. Ovarian steroidogenesis was also identified as a top disrupted pathways in two RNA-sequencing studies of adult mice exposed to PS nanoplastics (Xiong et al., 2024; Xue et al., 2024). Overall, the body of evidence on the disruption of ovarian steroidogenesis suggests that nanoplastics are endocrine disruptors, although, similar to classes of endocrine disrupting chemicals with variations in structure and properties like phthalates (Hannon and Flaws, 2015), the effects vary with properties such as size, shape, and polymer type.

We also surveyed expression of genes related to cell cycle, apoptosis, oxidative stress, and nuclear receptors in exposed follicles, which have been previously observed to be disrupted by endocrine disrupting chemicals and/or other nanoplastic exposures 39]. Each group of genes had one or two impacted genes that were mostly downregulated, but no clear pattern of effects emerged. Consistently, different genes were impacted by PS compared to PET. Future studies should expand this work using RNA-sequencing, ideally single-cell sequencing as antral follicles contain diverse cell types that we have previously shown are differentially impacted by endocrine disruptors (Mattson and Warner, 2025), to compared top disrupted pathways across various polymers, sizes, and shapes.

Cell cycle regulators play a critical role in follicle growth and have a key role in the process of ovarian folliculogenesis. Cell division is regulated by the cyclin-dependent kinase genes, which include genes tested in this experiment, including *Cdk4* and *Cdkn1a*, both of which were found to be downregulated due to PS NP exposure. Interestingly, *Cdk4* was found to be borderline upregulated in the second highest treatment group and then had a downregulation in the highest treatment group compared to control. Similar findings were observed in an *in vitro* study showing that PS microplastic exposure downregulates cyclin protein levels and results in cell cycle arrest in a mouse granulosa cell line, an effect that was exacerbated when combined with phthalate exposure (Wu et al., 2023). Since cell growth is essential for follicle maturation and oocyte support, disruption at the molecular level could lead to ovarian follicle development impairment (Balali et al., 2024). Moreover, our results show reduced follicle growth due to exposure, which could be related to cell cycle regulator disruption (Zhou and Flaws, 2017).

An interesting finding was the downregulation of *Ar* expression by PS in what appears to be a dose dependent manner. Androgen receptors are involved in regulating (and are regulated by) steroidogenesis and the cell cycle (Devillers et al., 2023; Koryakina et al., 2015) and may also be involved in regulation of apoptosis in granulosa cells (Risek et al., 2008). Specifically, decreased *Cdkn1a* may be related to decreased *Ar* from PS exposure.

Micro and nanoplastics have consistently been shown to induce oxidative stress in cells across tissue types (An et al., 2021; Ferrante et al., 2022) and contribute to inflammation, cellular damage, and disruption of normal physiological functions (Kadac-Czapska et al., 2024; Mahmud et al., 2024). In the ovary, oxidative stress disbalance can cause premature ovarian aging and may be related to disorders such as polycystic ovary syndrome (Yan et al., 2022). We measured levels of antioxidants and found *Sod1* was downregulated by a high dose of PS, similar to previous studies in antral follicles using phthalates (Wang et al., 2012; Zhou and Flaws, 2017). The downregulation of *Sod1* may indicate an increase in oxidative stress in the ovary.

In this work, we investigated two polymer types with distinct particle size and shapes. The smaller, perfectly spherical polystyrene nanoparticles disrupted different genes and hormones compared to the secondary PET nanoparticles synthesized in the lab, which were irregular in shape and slightly larger size than the PS nanoplastics. We chose the PET particles because they are a better representation of particles that humans would be exposed to in bottled water and food packaging from shedding plastic particles. However, because of changing multiple variables between the two plastic samples, we are not able to tell if the differing effects are caused by polymer type, shape, or size. We used lower doses for the PET particles to be more environmentally relevant and because the particle agglomerate at higher concentrations. Although we sonicated when the treatment solutions were prepared, the particles likely agglomerated to some extent in culture, which means that the follicles may have been exposed to the broader ranges of sizes indicated by the non-sonicated DLS analysis, including larger, although still nano-sized, clusters. In addition, the polymer types have different densities. PS particles are of low density and tend to float, which may have reduced exposure during follicle culture, whereas PET is high density and tends to sink, which would have increased follicle exposure (Forest and Pourchez, 2023).

In conclusion, we found that PS and PET nanoplastics at environmentally relevant doses caused disruption to cultured antral follicle function by reducing follicle growth and both increasing and decreasing hormone levels and expression of key genes related to important steroidogenic processes. The PS spheres and irregularly shaped secondary PET had different effects, although one was clearly more toxic than the other. The impacts on steroidogenesis indicate that the nanoplastics had endocrine disrupting effects and the impact on growth and the cell cycle suggest effects on folliculogenesis, although the mechanisms through which nanoplastics cause these effects are yet unknown. Further research into environmentally relevant doses of environmentally relevant plastics, as well as mixture studies, are necessary to improve our understanding of the potential health impacts of nanoplastics exposure.

## Data Availability Statement

Data will be made available by the corresponding author upon reasonable request.

## Acknowledgements

Thank you to all members of the EDC Lab at NJIT.

## Declaration of generative AI and AI-assisted technologies in the writing process

No generative AI technologies were used in the preparation of this manuscript.

## Author Contributions: Credit

**Hanin Alahmadi**: investigation, project administration, validation, visualization, writing - original draft, writing - reviewing and editing; **Maira Nadeem**: investigation, validation, writing original draft; **Alixs Pujols**: investigation, validation**; Raulle Reynolds**: investigation, validation; **Courtney Potts**: investigation, validation; **Allison Harbolic**: investigation; **Gania Lafontant**: visualization; **Indrani Gupta**: visualization; **Mohammad Saiful Islam**: visualization, validation; **Somenath Mitra**: resources, supervision, funding acquisition; **Genoa Warner**: conceptualization, methodology, validation, formal analysis, investigation, data curation, writing - original draft, writing - review and editing, visualization, supervision, project administration, funding acquisition.

## References

1. Afreen, V., Hashmi, K., Nasir, R., Saleem, A., Khan, M.I., Akhtar, M.F., 2023. Adverse health effects and mechanisms of microplastics on female reproductive system: a descriptive review. Environ Sci Pollut Res Int 30, 76283–76296. 10.1007/s11356-023-27930-1

2. An, R., Wang, X., Yang, L., Zhang, J., Wang, N., Xu, F., Hou, Y., Zhang, H., Zhang, L., 2021. Polystyrene microplastics cause granulosa cells apoptosis and fibrosis in ovary through oxidative stress in rats. Toxicology 449, 152665. 10.1016/j.tox.2020.152665

3. Annangi, B., Villacorta, A., Vela, L., Tavakolpournegari, A., Marcos, R., Hernández, A., 2023. Effects of true-to-life PET nanoplastics using primary human nasal epithelial cells. Environmental Toxicology and Pharmacology 100, 104140. 10.1016/j.etap.2023.104140

4. Avio, C.G., Gorbi, S., Regoli, F., 2017. Plastics and microplastics in the oceans: From emerging pollutants to emerged threat. Marine Environmental Research, Blue Growth and Marine Environmental Safety 128, 2–11. 10.1016/j.marenvres.2016.05.012

5. Balali, H., Morabbi, A., Karimian, M., 2024. Concerning influences of micro/nano plastics on female reproductive health: focusing on cellular and molecular pathways from animal models to human studies. Reproductive Biology and Endocrinology 22, 141. 10.1186/s12958-024-01314-7

6. Bashirova, N., Poppitz, D., Klüver, N., Scholz, S., Matysik, J., Alia, A., 2023. A mechanistic understanding of the effects of polyethylene terephthalate nanoplastics in the zebrafish (Danio rerio) embryo. Sci Rep 13, 1891. 10.1038/s41598-023-28712-y

7. Brits, M., van Velzen, M.J.M., Sefiloglu, F.Ö., Scibetta, L., Groenewoud, Q., Garcia-Vallejo, J.J., Vethaak, A.D., Brandsma, S.H., Lamoree, M.H., 2024. Quantitation of micro and nanoplastics in human blood by pyrolysis-gas chromatography–mass spectrometry. Microplastics and Nanoplastics 4, 12. 10.1186/s43591-024-00090-w

8. da Silva Brito, W.A., Mutter, F., Wende, K., Cecchini, A.L., Schmidt, A., Bekeschus, S., 2022. Consequences of nano and microplastic exposure in rodent models: the known and unknown. Particle and Fibre Toxicology 19, 28. 10.1186/s12989-022-00473-y

9. Devillers, M.M., François, C.M., Chester, M., Corre, R., Cluzet, V., Giton, F., Cohen-Tannoudji, J., Guigon, C.J., 2023. Androgen receptor signaling regulates follicular growth and steroidogenesis in interaction with gonadotropins in the ovary during mini-puberty in mice. Front Endocrinol (Lausanne) 14, 1130681. 10.3389/fendo.2023.1130681

10. Dubey, I., Khan, S., Kushwaha, S., 2022. Developmental and reproductive toxic effects of exposure to microplastics: A review of associated signaling pathways. Front Toxicol 4, 901798. 10.3389/ftox.2022.901798

11. Ferrante, M.C., Monnolo, A., Del Piano, F., Mattace Raso, G., Meli, R., 2022. The Pressing Issue of Micro- and Nanoplastic Contamination: Profiling the Reproductive Alterations Mediated by Oxidative Stress. Antioxidants (Basel) 11, 193. 10.3390/antiox11020193

12. Forest, V., Pourchez, J., 2023. Can the impact of micro- and nanoplastics on human health really be assessed using *in vitro* models? A review of methodological issues. Environment International 178, 108115. 10.1016/j.envint.2023.108115

13. Garcia, M.A., Liu, R., Nihart, A., El Hayek, E., Castillo, E., Barrozo, E.R., Suter, M.A., Bleske, B., Scott, J., Forsythe, K., Gonzalez-Estrella, J., Aagaard, K.M., Campen, M.J., 2024. Quantitation and identification of microplastics accumulation in human placental specimens using pyrolysis gas chromatography mass spectrometry. Toxicological Sciences 199, 81–88. 10.1093/toxsci/kfae021

14. Giri, S., Lamichhane, G., Khadka, D., Devkota, H.P., 2024. Microplastics contamination in food products: Occurrence, analytical techniques and potential impacts on human health. Current Research in Biotechnology 7, 100190. 10.1016/j.crbiot.2024.100190

15. González-Doncel, M., García-Mauriño, J.E., Beltrán, E.M., Fernández Torija, C., Andreu-Sánchez, O., Pablos, M.V., 2022. Effects of life cycle exposure to polystyrene microplastics on medaka fish (*Oryzias latipes*). Environmental Pollution 311, 120001. 10.1016/j.envpol.2022.120001

16. Haddadi, A., Kessabi, K., Boughammoura, S., Rhouma, M.B., Mlouka, R., Banni, M., Messaoudi, I., 2022. Exposure to microplastics leads to a defective ovarian function and change in cytoskeleton protein expression in rat. Environ Sci Pollut Res Int 29, 34594– 34606. 10.1007/s11356-021-18218-3

17. Hannon, P.R., Flaws, J.A., 2015. The effects of phthalates on the ovary. Frontiers in Endocrinology 6, 1–19. 10.3389/fendo.2015.00008

18. Hu, C.J., Garcia, M.A., Nihart, A., Liu, R., Yin, L., Adolphi, N., Gallego, D.F., Kang, H., Campen, M.J., Yu, X., 2024. Microplastic presence in dog and human testis and its potential association with sperm count and weights of testis and epididymis. Toxicological Sciences 200, 235–240. 10.1093/toxsci/kfae060

19. Huang, J., Zou, L., Bao, M., Feng, Q., Xia, W., Zhu, C., 2023. Toxicity of polystyrene nanoparticles for mouse ovary and cultured human granulosa cells. Ecotoxicology and Environmental Safety 249, 114371. 10.1016/j.ecoenv.2022.114371

20. Islam, M.S., Gupta, I., Xia, L., Pitchai, A., Shannahan, J., Mitra, S., 2024. Generation of Eroded Nanoplastics from Domestic Wastes and Their Impact on Macrophage Cell Viability and Gene Expression. Molecules 29, 2033. 10.3390/molecules29092033

21. Ji, Y., Wang, C., Wang, Y., Fu, L., Man, M., Chen, L., 2020. Realistic polyethylene terephthalate nanoplastics and the size- and surface coating-dependent toxicological impacts on zebrafish embryos. Environ. Sci.: Nano 7, 2313–2324. 10.1039/D0EN00464B

22. Kadac-Czapska, K., Ośko, J., Knez, E., Grembecka, M., 2024. Microplastics and Oxidative Stress—Current Problems and Prospects. Antioxidants (Basel) 13, 579. 10.3390/antiox13050579

23. Koelmans, A.A., Mohamed Nor, N.H., Hermsen, E., Kooi, M., Mintenig, S.M., De France, J., 2019. Microplastics in freshwaters and drinking water: Critical review and assessment of data quality. Water Research 155, 410–422. 10.1016/j.watres.2019.02.054

24. Koryakina, Y., Knudsen, K.E., Gioeli, D., 2015. Cell-cycle-dependent regulation of androgen receptor function. Endocrine-Related Cancer 22, 249–264. 10.1530/ERC-14-0549

25. Leslie, H.A., van Velzen, M.J.M., Brandsma, S.H., Vethaak, A.D., Garcia-Vallejo, J.J., Lamoree, M.H., 2022. Discovery and quantification of plastic particle pollution in human blood. Environment International 163, 107199. 10.1016/j.envint.2022.107199

26. Li, Y., Chen, L., Zhou, N., Chen, Y., Ling, Z., Xiang, P., 2024. Microplastics in the human body: A comprehensive review of exposure, distribution, migration mechanisms, and toxicity. Science of The Total Environment 946, 174215. 10.1016/j.scitotenv.2024.174215

27. Limonta, G., Mancia, A., Benkhalqui, A., Bertolucci, C., Abelli, L., Fossi, M.C., Panti, C., 2019. Microplastics induce transcriptional changes, immune response and behavioral alterations in adult zebrafish. Sci Rep 9, 15775. 10.1038/s41598-019-52292-5

28. Lin, X., Xie, H., Zhang, Y., Tian, X., Cui, L., Shi, N., Wang, L., Zhao, J., An, L., Wang, J., Li, B., Li, Y.-F., 2023. The toxicity of nano polyethylene terephthalate to mice: Intestinal obstruction, growth retardant, gut microbiota dysbiosis and lipid metabolism disorders. Food and Chemical Toxicology 172, 113585. 10.1016/j.fct.2022.113585

29. Liu, Z., Zhuan, Q., Zhang, L., Meng, L., Fu, X., Hou, Y., 2022. Polystyrene microplastics induced female reproductive toxicity in mice. Journal of Hazardous Materials 424, 127629. 10.1016/j.jhazmat.2021.127629

30. Mahmud, F., Sarker, D.B., Jocelyn, J.A., Sang, Q.-X.A., 2024. Molecular and Cellular Effects of Microplastics and Nanoplastics: Focus on Inflammation and Senescence. Cells 13, 1788. 10.3390/cells13211788

31. Manoochehri, Z., Etebari, M., Pannetier, P., Ebrahimpour, K., 2024. In vitro toxicity of polyethylene terephthalate nanoplastics (PET-NPs) in human hepatocarcinoma (HepG2) cell line. Toxicol. Environ. Health Sci. 16, 203–215. 10.1007/s13530-024-00213-z

32. Marcelino, R.C., Cardoso, R.M., Domingues, E.L.B.C., Gonçalves, R.V., Lima, G.D.A., Novaes, R.D., 2022. The emerging risk of microplastics and nanoplastics on the microstructure and function of reproductive organs in mammals: A systematic review of preclinical evidence. Life Sciences 295, 120404. 10.1016/j.lfs.2022.120404

33. Mason, S.A., Welch, V.G., Neratko, J., 2018. Synthetic Polymer Contamination in Bottled Water. Front. Chem. 6. 10.3389/fchem.2018.00407

34. Mattson, E., Warner, G.R., 2025. Single cell RNA-seq reveals that granulosa cells are a target of phthalate toxicity in the ovary. Toxicological Sciences kfaf001. 10.1093/toxsci/kfaf001

35. Montano, L., Raimondo, S., Piscopo, M., Ricciardi, M., Guglielmino, A., Chamayou, S., Gentile, R., Gentile, M., Rapisarda, P., Oliveri Conti, G., Ferrante, M., Motta, O., 2025. First evidence of microplastics in human ovarian follicular fluid: An emerging threat to female fertility. Ecotoxicol Environ Saf 291, 117868. 10.1016/j.ecoenv.2025.117868

36. Nayanathara Thathsarani Pilapitiya, P.G.C., Ratnayake, A.S., 2024. The world of plastic waste: A review. Cleaner Materials 11, 100220. 10.1016/j.clema.2024.100220

37. Nihart, A.J., Garcia, M.A., El Hayek, E., Liu, R., Olewine, M., Kingston, J.D., Castillo, E.F., Gullapalli, R.R., Howard, T., Bleske, B., Scott, J., Gonzalez-Estrella, J., Gross, J.M., Spilde, M., Adolphi, N.L., Gallego, D.F., Jarrell, H.S., Dvorscak, G., Zuluaga-Ruiz, M.E., West, A.B., Campen, M.J., 2025. Bioaccumulation of microplastics in decedent human brains. Nat Med 1–6. 10.1038/s41591-024-03453-1

38. Patel, S., Zhou, C., Rattan, S., Flaws, J.A., 2015. Effects of Endocrine-Disrupting Chemicals on the Ovary1. Biology of Reproduction 93, 20, 1–9. 10.1095/biolreprod.115.130336

39. Pfaffl, M.W., 2001. A new mathematical model for relative quantification in real-time RT–PCR. Nucleic Acids Res 29, e45.

40. Qian, N., Gao, X., Lang, X., Deng, H., Bratu, T.M., Chen, Q., Stapleton, P., Yan, B., Min, W., 2024. Rapid single-particle chemical imaging of nanoplastics by SRS microscopy. Proceedings of the National Academy of Sciences 121, e2300582121. 10.1073/pnas.2300582121

41. Risek, B., Bilski, P., Rice, A.B., Schrader, W.T., 2008. Androgen Receptor-Mediated Apoptosis Is Regulated by Photoactivatable Androgen Receptor Ligands. Molecular Endocrinology 22, 2099–2115. 10.1210/me.2007-0426

42. Senathirajah, K., Attwood, S., Bhagwat, G., Carbery, M., Wilson, S., Palanisami, T., 2021. Estimation of the mass of microplastics ingested – A pivotal first step towards human health risk assessment. Journal of Hazardous Materials 404. 10.1016/j.jhazmat.2020.124004

43. Sussarellu, R., Suquet, M., Thomas, Y., Lambert, C., Fabioux, C., Pernet, M.E.J., Le Goïc, N., Quillien, V., Mingant, C., Epelboin, Y., Corporeau, C., Guyomarch, J., Robbens, J., Paul-Pont, I., Soudant, P., Huvet, A., 2016. Oyster reproduction is affected by exposure to polystyrene microplastics. Proceedings of the National Academy of Sciences 113, 2430– 2435. 10.1073/pnas.1519019113

44. Wang, Q., Chi, F., Liu, Y., Chang, Q., Chen, S., Kong, P., Yang, W., Liu, W., Teng, X., Zhao, Y., Guo, Y., 2025. Polyethylene microplastic exposure adversely affects oocyte quality in human and mouse. Environment International 195, 109236. 10.1016/j.envint.2024.109236

45. Wang, W., Craig, Z.R., Basavarajappa, M.S., Hafner, K.S., Flaws, J.A., 2012. Mono-(2- Ethylhexyl) Phthalate Induces Oxidative Stress and Inhibits Growth of Mouse Ovarian Antral Follicles. Biology of Reproduction 87, 1–10. 10.1095/biolreprod.112.102467

46. Wei, Z., Wang, Y., Wang, S., Xie, J., Han, Q., Chen, M., 2022. Comparing the effects of polystyrene microplastics exposure on reproduction and fertility in male and female mice. Toxicology 465, 153059. 10.1016/j.tox.2021.153059

47. Winkler, A., Santo, N., Ortenzi, M.A., Bolzoni, E., Bacchetta, R., Tremolada, P., 2019. Does mechanical stress cause microplastic release from plastic water bottles? Water Research 166, 115082. 10.1016/j.watres.2019.115082

48. Wu, H., Liu, Q., Yang, N., Xu, S., 2023. Polystyrene-microplastics and DEHP co-exposure induced DNA damage, cell cycle arrest and necroptosis of ovarian granulosa cells in mice by promoting ROS production. Science of The Total Environment 871, 161962. 10.1016/j.scitotenv.2023.161962

49. Xiong, G., Zhang, H., Peng, Y., Shi, H., Han, M., Hu, T., Wang, H., Zhang, S., Wu, X., Xu, G., Zhang, J., Liu, Y., 2024. Subchronic co-exposure of polystyrene nanoplastics and 3-BHA significantly aggravated the reproductive toxicity of ovaries and uterus in female mice. Environmental Pollution 351, 124101. 10.1016/j.envpol.2024.124101

50. Xue, Y., Cheng, X., Ma, Z.-Q., Wang, H.-P., Zhou, C., Li, J., Zhang, D.-L., Hu, L.-L., Cui, Y.- F., Huang, J., Luo, T., Zheng, L.-P., 2024. Polystyrene nanoplastics induce apoptosis, autophagy, and steroidogenesis disruption in granulosa cells to reduce oocyte quality and fertility by inhibiting the PI3K/AKT pathway in female mice. J Nanobiotechnol 22, 460. 10.1186/s12951-024-02735-7

51. Yan, F., Zhao, Q., Li, Y., Zheng, Z., Kong, X., Shu, C., Liu, Y., Shi, Y., 2022. The role of oxidative stress in ovarian aging: a review. Journal of Ovarian Research 15, 100. 10.1186/s13048-022-01032-x

52. Yang, J., Kamstra, J., Legler, J., Aardema, H., n.d. The impact of microplastics on female reproduction and early life. Anim Reprod 20, e20230037. 10.1590/1984-3143-AR2023-0037

53. Zarus, G.M., Muianga, C., Hunter, C.M., Pappas, R.S., 2021. A review of data for quantifying human exposures to micro and nanoplastics and potential health risks. Science of The Total Environment 756, 144010. 10.1016/j.scitotenv.2020.144010

54. Zeng, L., Zhou, C., Xu, W., Huang, Y., Wang, W., Ma, Z., Huang, J., Li, J., Hu, L., Xue, Y., Luo, T., Zheng, L., 2023. The ovarian-related effects of polystyrene nanoplastics on human ovarian granulosa cells and female mice. Ecotoxicology and Environmental Safety 257, 114941. 10.1016/j.ecoenv.2023.114941

55. Zhang, H., Zhang, S., Duan, Z., Wang, L., 2022. Pulmonary toxicology assessment of polyethylene terephthalate nanoplastic particles *in vitro*. Environment International 162, 107177. 10.1016/j.envint.2022.107177

56. Zhang, J., Wang, L., Kannan, K., 2020. Microplastics in house dust from 12 countries and associated human exposure. Environment International 134, 105314. 10.1016/j.envint.2019.105314

57. Zhang, J., Wang, L., Trasande, L., Kannan, K., 2021. Occurrence of Polyethylene Terephthalate and Polycarbonate Microplastics in Infant and Adult Feces. Environmental Science and Technology Letters 8, 989–994. 10.1021/acs.estlett.1c00559

58. Zhang, J., Zou, Y., Hu, L., Zhao, Y., Fen, Y., Xu, H., 2023. TiO2 nanoparticles combined with polystyrene nanoplastics aggravated reproductive toxicity in female mice via exacerbating intestinal barrier disruption. J Sci Food Agric. 10.1002/jsfa.12722

59. Zhang, Y., Hales, B.F., Robaire, B., 2025. Exposure to polystyrene nanoplastics induces lysosomal enlargement and lipid droplet accumulation in KGN human ovarian granulosa cells. Arch Toxicol 99, 1445–1454. 10.1007/s00204-025-03969-6

60. Zhou, C., Flaws, J.A., 2017. Effects of an environmentally relevant phthalate mixture on cultured mouse antral follicles. Toxicological Sciences 156, 217–229. 10.1093/toxsci/kfw245

61. Zuccarello, P., Ferrante, M., Cristaldi, A., Copat, C., Grasso, A., Sangregorio, D., Fiore, M., Oliveri Conti, G., 2019. Exposure to microplastics (<10 μm) associated to plastic bottles mineral water consumption: The first quantitative study. Water Research 157, 365–371. 10.1016/j.watres.2019.03.091

