## Supplementary Figure for "Polystyrene and polyethylene terephthalate nanoplastics differentially impact mouse ovarian follicle function"

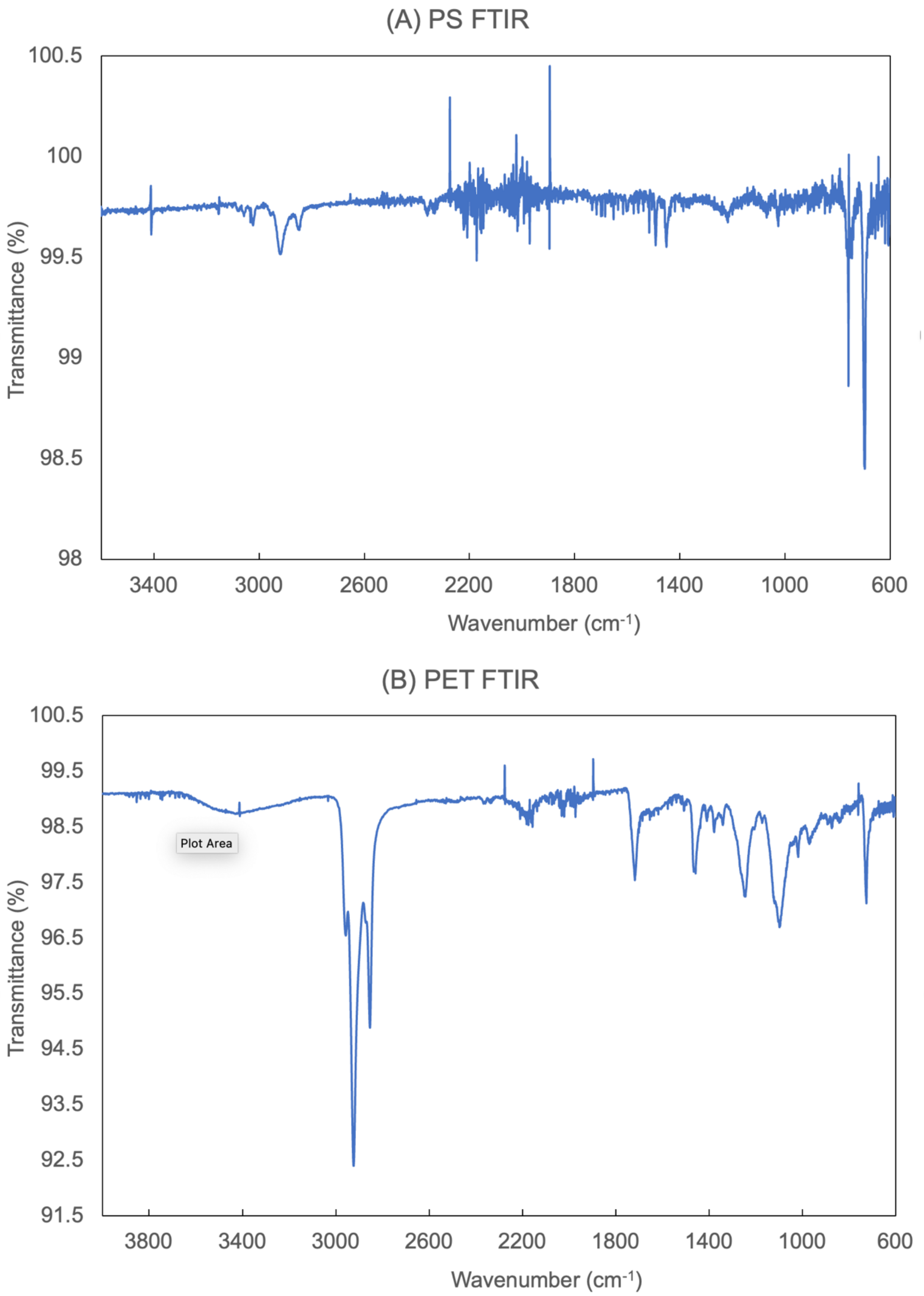


**Figure S1** Polymer type was confirmed using Fourtier transform infrared spectroscopy (FTIR) for PS (A) and PET (B).
